# Kaposi Sarcoma Herpes Virus Reprograms Mesenchymal Cell Glycosylation to control platelet-derived growth factor receptor A signaling

**DOI:** 10.64898/2026.06.29.735238

**Authors:** Julián Gambarte Tudela, Patricia N. Gutiérrez, M.A. Montani, Nadia Bannoud, Pablo A. García, Karina V. Mariño, Martin Abba, Juan P. Cerliani, Julián Naipauer, Ezequiel Lacunza, Enrique A Mesri, Gabirel A. Rabinovich, Diego O. Croci

**Affiliations:** Instituto de Histología y Embriología de Mendoza (IHEM), Consejo Nacional de Investigaciones Científicas y Técnicas (CONICET), Universidad Nacional de Cuyo. M5500, Mendoza, Argentina; UM-CFAR/SCCC Argentina Consortium for Research and Training in Virally Induced AIDS-Malignancies, Miami, USA; Instituto de Fisiología, Biología Molecular y Neurociencias (IFIBYNE), CONICET-Universidad de Buenos Aires, Buenos Aires, Argentina; Facultad de Ciencias Médicas, Universidad Nacional de Cuyo. M5500, Mendoza, Argentina; Laboratorio de Glicómica Funcional y Molecular, Instituto de Biología y Medicina Experimental (IBYME), Consejo Nacional de Investigaciones Científicas y Técnicas (CONICET), C1428ADN Buenos Aires, Argentina; Universidad Argentina de la Empresa (UADE), Instituto de Tecnología (INTEC), C1073 Ciudad de Buenos Aires, Argentina; Centro de Investigaciones Inmunológicas Básicas y Aplicadas, Facultad de Ciencias Médicas, Universidad Nacional de La Plata, La Plata, Argentina; Laboratorio de Glicomedicina, Instituto de Biología y Medicina Experimental (IBYME), Consejo Nacional de Investigaciones Científicas y Técnicas (CONICET), C1428ADN Buenos Aires, Argentina; Laboratory of Glycoimmunology, CaixaResearch Institute, 08035, Barcelona, Spain; Facultad de Ciencias Exactas y Naturales, Universidad Nacional de Buenos Aires, C1428 Ciudad de Buenos Aires, Argentina; Instituto Tecnológico de Buenos Aires, ITBA. C1437ECC Buenos Aires, Argentina

## Abstract

Kaposi sarcoma–associated herpesvirus (KSHV) reprograms host cellular pathways to promote viral persistence and tumor development by supporting the survival and expansion of infected and bystander cells. Among the signaling axes implicated in this process, oncogenic activation of receptor tyrosine kinases, particularly platelet-derived growth factor receptor alpha (PDGFRA), is a well-established hallmark of Kaposi sarcoma pathogenesis. However, the impact of virus-driven cell surface glycosylation changes in PDGFRA-associated signaling remains uncertain.

Here, we identified a critical role for KSHV in reshaping the glycosylation landscape of mesenchymal stromal cells (MSCs), thereby reprogramming PDGFRA signaling. We show that KSHV infection enhances a unique glycan profile in human and mouse MSCs, enriched in branched complex N-glycans with limited α(2-6) sialylation, Transcriptomic analyses of KSHV-infected MSCs and Kaposi’s Sarcoma patient samples revealed a coordinated dysregulation of pathways involved in carbohydrate metabolism, nucleotide-sugar transport, and sialic acid turnover, with a particular focus on sialic acid turnover (NPL/NEU3), the UDP-N-acetylglucosamine transporters SLC35A3/B4, and complex N-glycan elongation (MGAT5/B3GNT2/B4GALT1) as critical nodes underlying reduced α(2-6) sialylation and increased N-glycan branching.

This remodeled glycan landscape contributes to create a permissive context for galectin-1 (Gal-1) binding, which in turn enhances PDGFRA activation and downstream signaling. These findings identify an integral component of KSHV-driven mesenchymal cell reprogramming, unveiling a lectin-dependent mechanism of RTK activation in Kaposi sarcoma pathogenesis.

## Introduction

Kaposi’s sarcoma–associated herpesvirus (KSHV) is the etiological agent of Kaposi’s sarcoma (KS), a prominent acquired immunodeficient syndrome (AIDS)-defining malignancy ^1,2^. KSHV represents a paradigmatic example of a tumorigenic virus that hijacks host signaling networks without introducing fundamentally new biological programs ^3,4^. Instead, KSHV exploits pre-existing cellular pathways involved in vascular growth, inflammation and stromal plasticity, promoting sustained activation states that are permissive for tumor persistence ^2,4^. Among these pathways, receptor tyrosine kinase (RTK) signaling has emerged as a central node in KS pathogenesis, particularly within mesenchymal and stromal compartments of the tumor microenvironment (TME) ^3,5^.

Platelet-derived growth factor receptor-α (PDGFRA), a critical signaling RTK, has emerged as a dominant effector of KSHV-driven sarcomagenesis. Genetic, biochemical, and *in vivo* studies have shown that KSHV, largely through its viral G protein-coupled receptor (vGPCR), promotes PDGFRA activation and sustains proliferative and angiogenic programs in mesenchymal-like infected cells ^6–9^. MSCs are particularly relevant in this context, as they display marked susceptibility to KSHV-induced reprogramming and have been proposed as biologically relevant cell targets of KS pathogenesis ^5^

Beyond ligand availability and receptor expression levels, the cell surface glycosylation landscape has emerged as a critical determinant of growth factor receptor signaling, altering its organization, membrane retention, and activation threshold independently of canonical inputs^10–12^. Within this regulatory framework, galectins, a family of soluble β-galactoside–binding proteins, have emerged as direct transducers of glycan information into functional signaling outputs, bridging glycan composition and receptor activation state^11,13,14^.

Galectin-1 (Gal-1; encoded by *LGALS1* gene) has been implicated in the regulation of vascular inflammation and tumor signaling through its ability to engage glycosylated cell surface receptors ^15–19^. In previous studies, we demonstrated that Gal-1 can modulate the threshold of vascular endothelial growth factor (VEGF) receptor 2 signaling through a glycosylation-dependent mechanism. This lectin functions as a non-canonical ligand, sustaining receptor activation independently of VEGF ^20^. This glycosylation-dependent regulatory layer has emerged as a key mechanism underlying persistent receptor signaling in pathological settings, including cancer, fibrosis and chronic inflammation^21^.

Despite extensive characterization of vGPCR-driven signaling in KS oncogenesis ^8,22,23^ and growing recognition of glycosylation-dependent RTK regulation in cancer ^20,24–27^, whether KSHV reshapes the glycosylation landscape of mesenchymal cells, and whether such remodeling influences lectin-mediated RTK signaling has not been addressed.

Here, we show that KSHV infection and vGPCR expression coordinately remodel the glycosylation profile of mesenchymal stromal cells, creating a permissive context for Gal-1 engagement and PDGFRA activation. By integrating lectin-based glycophenotyping, transcriptomic analyses, and functional studies of receptor signaling and trafficking, here we identify a viral-driven glycosylation program that enables Gal-1–dependent PDGFRA activation. These findings reveal a previously unrecognized layer of host–virus interaction, highlighting glycosylation remodeling as a critical regulator of receptor signaling in viral-driven oncogenesis.

## Materials and Methods

### Cells and Reagents

Mesenchymal stromal cells (MSCs) were used as an experimental model based on their proposed role in KS pathogenesis ^5,28,29^. Mouse bone marrow–derived MSCs (BM-MSCs) were isolated from femurs of C57BL/6 mice as previously described ^30^ and maintained in α-MEM supplemented with 15% fetal bovine serum (FBS; Sigma) and antibiotic-antimycotic solution (Gibco). Cells were used at passage ≤5. Human umbilical cord–derived MSCs (hMSCs) were obtained and characterized as previously described ^31^. The murine bone marrow stromal cell line OP9 (ATCC CRL-2749) was cultured in DMEM GlutaMAX™ (Gibco) supplemented with 10% FBS (Sigma). iSLK.KSHV219 cells (human clear cell renal carcinoma, male; RRID: CVCL_9569) were cultured in DMEM (Gibco) supplemented with 10% FBS (Sigma) and 1% penicillin-streptomycin (Gibco) at 37°C in 5% CO₂, as previously described ^32^, and maintained under selection with puromycin (10 μg/mL; Gibco), G418 (50 μg/mL; Sigma), and hygromycin B (50 μg/mL; Invitrogen).

Recombinant cytokines and growth factors used in this study included rhIL-6 (cat. 7270-IL), PDGF-AA (cat. 221-AA), PDGF-BB (cat. 220-BB), and VEGF (cat. 293-VE), all from R&D Systems; and FGF-2 (cat. SRP4037; Sigma-Aldrich). Lectins were used unconjugated and included: *Sambucus nigra* agglutinin (SNA), *Phaseolus vulgaris* leukoagglutinin (PHA-L), *Lycopersicon esculentum* lectin (LEL), Concanavalin A (ConA), peanut agglutinin (PNA), *Maackia amurensis* lectin (MAA), *Erythrina cristagalli* lectin (ECL), and *Aleuria aurantia* lectin (AAL), all from Vector Labs.

### Viral Constructs, Infection, and Transfection

iSLK.KSHV219 cells harboring recombinant KSHV.219 were used for virus preparation^33^. Briefly, infectious virus particles of the 219 strain were induced from the respective iSLK cells by treatment with doxycycline and sodium butyrate for 4 days. The culture supernatants were filtered through a 0.45 μm filter and centrifuged at 300 x g for 5 minutes. Cell-free virus-containing supernatants were used for further infections.

A synthetic vGPCR coding sequence (VGPCR_HHV8P, 1029 bp) was cloned into the pUC57 vector (GenScript). BM-MSCs and hMSCs were seeded and cultured until they reached approximately 80% confluence. Cells were washed once with serum-free α-MEM and transfected with 0.5 µg of vGPCR-GFP plasmid using X-treme Gene-9 transfection reagent (Merck) according to the manufacturer’s instructions.

Twenty hours after transfection, cells were washed twice with Tris-buffered saline (TBS) to remove dead cells and debris and subsequently maintained in α-MEM under standard culture conditions (37°C, 5% CO₂, 95% humidified air) until further analyses.

### Confocal microscopy

MSCs were transfected with either vGPCR or GFP (control) and seeded on 12-mm glass coverslips. After the indicated treatment, cells were fixed with 4% paraformaldehyde for 10 min at room temperature (RT) and washed three times with PBS. Cells were then incubated for 1 h. at RT with FITC-conjugated anti-Gal-1 monoclonal antibody (mAb)^20^ or with fluorescently labeled lectins SNA and L-PHA (Vector Labs). After three washes with PBS, the nuclei were stained with DAPI for 10 min at RT under gentle agitation. Images were acquired using an Olympus FV-1000 laser confocal microscope, and fluorescence intensity of Gal-1 and lectin binding was quantified using FIJI software.

MSCs were incubated with recombinant Gal-1 (3 μM) for 0, 5, 15, 30, or 45 min. Positive controls included PDGF-BB (5 nM) or 10% FCS. At each time point, cells were fixed with 4% paraformaldehyde and stained for phospho-PDGFRA (Tyr754, clone 23B2, Cell Signaling Technology) for 1 h. at RT, followed by incubation with an Alexa Fluor 555 conjugated secondary antibody (Cell Signaling Technology) for 1 h. Nuclei were counterstained with DAPI when indicated.

### Flow cytometry and lectin profiling

#### Lectin binding and glycophenotyping

MSCs were incubated with Zombie Green viability dye (BioLegend) according to manufacturer’s instructions. Cells were washed and incubated with fluorescently labeled lectins for 45 min at 4°C in the lectin buffer (150 mM NaCl, 10 mM HEPES, 1% BSA). The relative median fluorescence intensity (rMFI) was calculated as a ratio of the median fluorescence intensity of cells of interest to the median mean fluorescence intensity of negative control cells.

Lectins used included L-PHA, LEL, SNA, MAL II, PNA, ECL, and AAL all from (Vector Labs). Lectins were fluorescently labeled using the Lightning-Link Fast conjugation Kit (Abcam) according to the manufacturer’s instructions. For lectin binding assays, Dylight-488-labeled Gal-1, Gal-2 or Gal-7 were incubated with MSCs transfected with vGPCR vector or infected with KSHV for 1 h. at 4°C.

#### Intracellular phospho-protein detection (phospho-flow)

Phosphorylated STAT3 (Tyr705) and HIF-1α levels were assessed using a phospho-flow protocol (BD Biosciences, San Jose, CA, USA). MSCs incubated or not with recombinant Gal-1 were fixed with 2% paraformaldehyde (PFA) for 10 min at RT and permeabilized with BD Phosflow Perm Buffer III for 30 min on ice, according to the manufacturer’s instructions. Cells were washed with BD Phosflow Perm/Wash Buffer I and incubated with PE-conjugated anti-pSTAT3 (Tyr705 clone 4/P-STAT3) antibody at a 1:100 dilution for 45 min at 4 °C in the dark. Cells were stained with Alexa Fluor 647-conjugated mouse anti-HIF-1α antibody (BioLegend) at a 1:200 dilution in Phosflow Perm/Wash Buffer I. After staining, cells were washed twice with PBS containing 1% BSA and resuspended in PBS for acquisition. HIF-1α and pSTAT3 levels were quantified as rMFI within gated cell populations.

### MSC phenotyping

Human and murine MSCs were phenotypically characterized by flow cytometry to confirm identity and purity prior to downstream functional analyses. Human MSCs were analyzed using the BD Stemflow™ Human MSC Analysis Kit (BD Biosciences, Cat. No. 562245), which detects the minimal panel of surface markers recommended by the International Society for Cellular Therapy (ISCT), including positive and negative markers. Human MSCs were defined as ≥95% positive for CD105, CD73, and CD90 and ≤2% positive for hematopoietic lineage markers. For murine MSCs, a comparable panel of fluorochrome-conjugated antibodies recognizing the corresponding murine markers was used following an analogous staining and acquisition protocol. Data were acquired using a FACSAaria III or an Accury C6 Plus flow cytometer (Becton Dickinson) and analyzed with FlowJo software (version 10.7.1; FlowJo, LLC).

### Western blot analysis

Cells were lysed in lysis buffer containing 50 mM Tris-HCl (pH 7.5), 150 mM NaCl, 10 mM EDTA, and 1% NP-40, supplemented with protease and phosphatase inhibitor cocktails (Sigma-Aldrich, St. Louis, MO, USA). Protein concentration was determined using the MicroBCA Protein Assay Kit (Pierce, Thermo Fisher Scientific) according to the manufacturer’s instructions. Equal amounts of protein were mixed with Laemmli sample buffer (Bio-Rad, Hercules, CA, USA), resolved by SDS-PAGE, and transferred onto nitrocellulose membranes (GE Healthcare, Chicago, IL, USA). Membranes were blocked with 5% BSA in TBS and incubated overnight at 4°C with gentle agitation with primary antibodies at the following dilutions: anti-Erk1/2 (1/1000), anti-phospho-Erk1/2 (1/1000), anti-actin (1/4000), (all from Santa Cruz Biotechnology, Dallas, TX, USA); anti-STAT-3 (1/1000), anti-Phospho-STAT-3 (1/1000), anti-Akt (1/1000), anti-phospho-Akt (1/1000), anti-PDGFRA (1/100), anti-phospho-PDGFRA (1/100) (all from Cell Signaling Technology, Danvers, MA, USA) (all from Abcam, Cambridge, UK) or a polyclonal rabbit anti-Gal1 IgG (1.5 μg/mL)^20^.

For experiments analyzing phosphorylated and total proteins, membranes were first cut into regions corresponding to proteins of interest and incubated with phospho-specific antibodies. Membranes were stripped using (1.5% Glycine, 1% SDS, 0.1% Tween-20, pH 2.2) stripping buffer for 30 minutes, re-blocked with 5% BSA in TBS and re-probed overnight at 4°C with antibodies against the corresponding total proteins. After each primary antibody incubation, membranes were washed and incubated for 1 h at room temperature with HRP-conjugated secondary antibodies (Bio-Rad) under gentle agitation. Blots were developed using Immobilon chemiluminescent HRP substrate (Millipore, Burlington, MA, USA), and images were acquired using ImageQuant LAS 4000 (GE Healthcare). Band intensities were quantified using ImageJ software (version 1.54p, NIH, Bethesda, MD, USA).

### Transcriptomic Data Acquisition and Processing

Raw RNA-seq data were obtained from publicly available studies and prior work from our UM-CFAR/SCCC-Argentina consortium to investigate transcriptional changes associated with KSHV infection. Transcriptomic data from KSHV-infected human MSCs was generated in a previously described model^28^ (GSE260925) and analyzed together with an integrative KS transcriptomic dataset^34^. This dataset combines newly generated RNA-seq data from nine KS lesions (GSE271303) with three publicly available cohorts (GSE147704, SRP486827, and GSE241095), resulting in a harmonized dataset comprising 131 samples, including control skin and KS tumors stratified into previously defined molecular clusters. All datasets were processed using a unified analysis pipeline. Differential expression analyses were performed by comparing KS tumors with control skin samples in the integrative KS cohort and KSHV-infected MSCs with uninfected controls in the *in vitro* infection model. Differential gene expression was assessed using DESeq2 implemented in R through the Bioconductor framework. Functional enrichment analyses were performed with cluster Profiler, focusing on biological processes related to carbohydrate metabolism and glycosylation.

For visualization of gene expression patterns, heatmaps and violin plots were generated using the *pheatmap* and *vioplot* packages in R, respectively.

Differential expression analysis of glycosylation-related genes in KS lesions compared with control skin, grouped by functional category, was additionally performed using the UCSC Xena Browser, leveraging a public Xena Hub hosting batch-corrected gene expression and associated phenotypic data from KS cohorts (https://kaposi.xenahubs.net/).

### Glycosylation-related Gene Set Curation

To interrogate glycosylation-associated transcriptional programs, we assembled a curated set of genes implicated in glycan biosynthesis, modification, and recognition (‘glycogenes’). Gene selection was anchored in comprehensive glycosylation pathway atlases, including the human glycan biosynthetic maps described ^35,36^, which organize glycosyltransferases into defined functional modules corresponding to distinct glycan classes (e.g., N-linked, O-linked, glycolipid, and glycosaminoglycan pathways). These atlases provide a structured framework for identifying genes whose products directly contribute to glycan assembly and structural diversification.

This initial set was expanded to include additional functional categories relevant to glycan metabolism, including glycosidases, enzymes involved in the synthesis and interconversion of nucleotide sugar donors, Golgi nucleotide-sugar transporters, and glycan-binding proteins that interpret glycan structures. The final curated gene set encompasses enzymes, transporters, and receptors directly associated with glycan production and recognition.

This glycogene set served as the basis for downstream differential expression analysis and functional enrichment studies in transcriptomic datasets derived from KSHV-infected cells and Kaposi’s sarcoma samples, as described in the following section.

### Statistical Analysis

Statistical analysis was performed using GraphPad Prism 10 Software (GraphPad Software, Boston, MA, USA) or R software (version 4.4.2, R core team, 2024). Data represent the mean ± SEM of N independent experiments. For comparisons between two groups, an unpaired Student’s t-test was applied. For multiple comparisons, one-way or two-way ANOVA was used, followed by Bonferroni, Dunnett, or Tukey’s post-hoc tests, as appropriate. *P* values <0.05 were considered statistically significant.

## RESULTS

### Viral G protein-coupled receptor (vGPCR) expression remodels MSC glycosylation enhancing Gal-1 binding

To determine whether KSHV-driven signaling influences the glycosylation landscape of MSCs, we analyzed the cell surface glycophenotype of BM-MSCs under basal conditions and following vGPCR expression. BM-MSCs purity and identity were confirmed by flow cytometry (Figure S1A), and cell surface glycan structures were profiled using a multicolor lectin-based flow cytometry approach^37^, with a panel of fluorochrome-conjugated lectins selected to inform glycan motifs relevant for galectin binding (Figure S1B).

Gal-1 recognizes N-acetyllactosamine (LacNAc; Galβ1-4GlcNAc) units displayed in N- or O-linked glycans present in cell surface glycosylated receptors^14^. Terminal α(2-6) sialylation of LacNAc structures inhibits Gal-1 binding, whereas α(2-3) sialylation remains compatible with lectin engagement^38^. This biochemical framework provided the basis for assessing whether MSCs display a Gal-1-permissive glycosylation profile.

Under basal conditions, BM-MSCs displayed a glycosylation profile consistent with Gal-1: low reactivity to the *Sambuccus nigra* agglutinin (SNA), indicating limited α(2-6) sialylation, while consistent binding of *Maackia amurensis* agglutinin (MAL)-II, confirmed the presence of α(2-3)-sialylated structures. Moreover, these cells showed lower frequency of terminal galactose-containing epitopes detected by *Erythrina cristagalli* (ECL) And robust L-phytohemagglutinin (L-PHA) and *Lycopersicon esculentum* (LEL) staining, showing abundant β(1-6)–branched N-glycans and poly-LacNAc extension. Finally, detectable concanavalin A (ConA) and *Aleuria aurantia* lectin (AAL) reactivity indicated the presence of high-mannose N-glycans and core fucosylation (Figure 1A and 1B). Collectively these data support a poly-LacNAc and complex branched N-glycan surface landscape highly permissive for Gal-1 binding. Furthermore, assessment of O-glycan structures revealed a modest presence of peanut (PNA)–reactive asialo core-1 O-glycans (Figure 1A and 1B), which could serve as precursors for LacNAc-elongated core-2 structures required for Gal-1 binding (Figure S1B).

**Figure 1.**
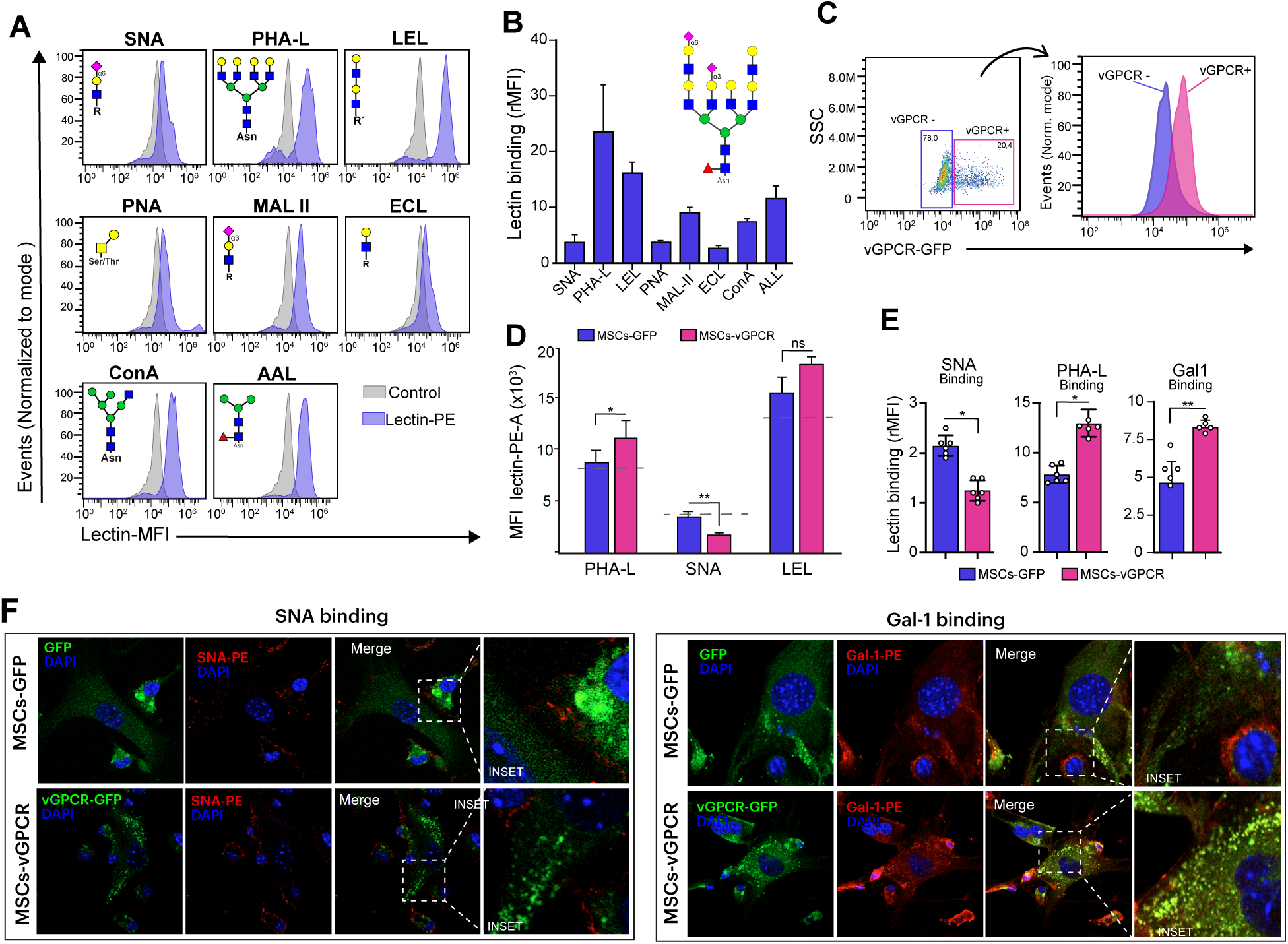
Glycophenotypic characterization and downstream analyses of vGPCR-expressing BM-MSCs. Representative lectin-based glycophenotypic profiling of mouse MSCs using SNA, L-PHA, LEL, ConA, PNA, MAL-II, ECL, and AAL analyzed by flow cytometry. B) Quantification of lectin binding shown in A, expressed as relative mean fluorescence intensity rMFI= ((MFI with lectin – MFI without lectin) / MFI without lectin. C) Flow cytometry analysis of mouse MSC transfected with vGPCR-GFP expression vector, including gating strategy to segregate vGPCR (+) and vGPCR (−) cells. Representative of three independent experiments. D) Comparative lectin-binding analysis of vGPCR (+) and vGPCR (-) bystander mouse MSCs using L-PHA, SNA, and LEL, quantified as rMFI. E) Flow cytometry analysis of SNA, L-PHA, and Gal-1 binding in mMSCs transfected with vGPCR or GFP empty vector. F) Confocal microscopy images of SNA and Gal-1 binding in vGPCR-GFP and empty-GFP transfected mouse MSCs; nuclei counterstained with DAPI. Images are representative of four independent experiments. Barrs represents the mean ± SEM; **P<0.01, *P<0.05.

Expression of vGPCR in BM-MSCs induced marked remodeling of this glycophenotypic profile (Figure 1C and 1D). Specifically focused on N-glycan structures, vGPCR-expressing cells showed reduced SNA binding and increased L-PHA and LEL reactivity, indicating enhanced exposure of branched poly-LacNAc containing complex N-glycans (Figure 1D); these glycosylated, structures favor GAL1 cross-linking.

To determine whether these glycosylation changes translated into altered galectin binding, we assessed Gal-1, L-PHA and SNA binding by confocal microscopy. Consistent with the flow cytometry data, vGPCR-expressing BM-MSCs showed increased L-PHA and reduced SNA binding (Figure 1E). Importantly, fluorescently labeled Gal-1 displayed significantly enhanced binding to vGPCR-expressing cells compared with controls, with the lectin predominantly retained at the cell surface rather than internalized (Figure 1E-F and S1C), indicating that vGPCR signaling is sufficient to activate a regulatory context permissive for Gal-1 binding.

Thus, BM-MSCs basally display a Gal-1–permissive glycan landscape, and vGPCR expression coordinately remodels this profile, reducing α(2-6) sialylation and expanding complex branched poly-LacNAc-rich N-glycan structures that enhance Gal-1 binding and signaling.

### KSHV-driven remodeling of cell surface glycosylation enhances Gal-1 binding to human MSCs

Having established that vGPCR expression remodels the glycosylation landscape of murine MSCs, we next examined whether these features are recapitulated in human MSCs (hMSCs) upon infection with a recombinant virus containing the complete KSHV genome (rKSHV.219)^33^. hMSCs were isolated from umbilical cord perivascular tissue and phenotypically characterized by flow cytometry (Figure S2A and S2B).

Under basal conditions, hMSCs displayed a cell surface glycoprofile compatible with Gal-1 engagement, evidenced by strong binding of LEL and PHA-L, and low SNA reactivity (Figure 2A), consistent with β(1,6) branched N-glycans containing multiple LacNAc units and low α(2,6) sialylation. Fluorescently labeled Gal-1 readily bound to human MSCs under these conditions. Interestingly, other prototypic galectins with different sialylation restrictions, implicated in epithelial and skin inflammatory responses, including Gal-2 and Gal-7, displayed comparable basal binding intensities (Figure 2B), indicating that the Gal-1–permissive glycan landscape of hMSCs is not lectin-specific under resting conditions.

**Figure 2.**
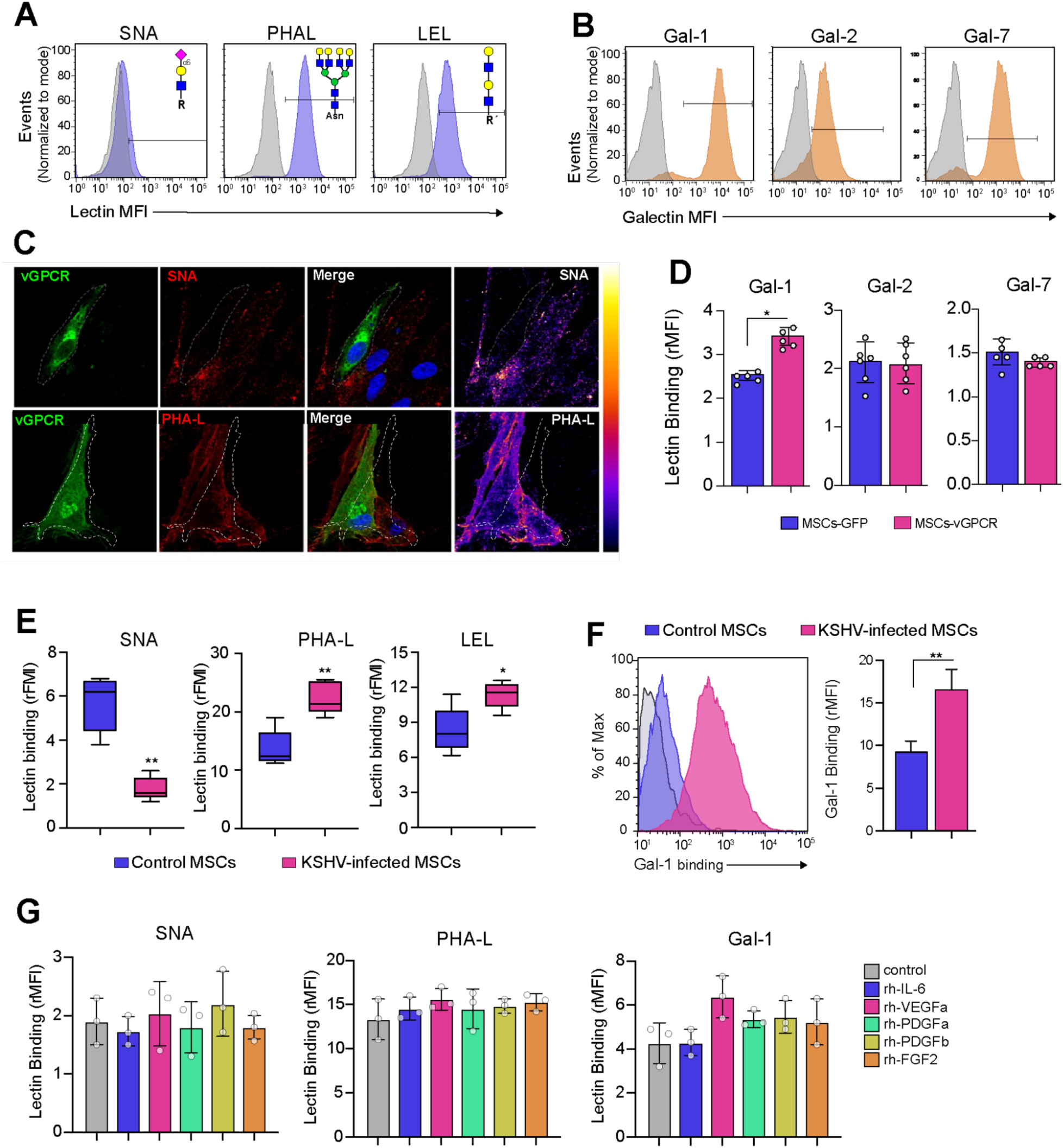
KSHV infection and vGPCR expression selectively enhance Gal-1 engagement through glycosylation remodeling in human MSCs. A) Flow cytometry Lectin-based glycophenotypic profiling of human MSCs using SNA, PHA-L, and LEL. Histograms are representative of five independent experiments. B) Flow cytometry analysis of recombinant Gal-1, Gal-2, and Gal-7 binding to human MSCs. Histograms are representative of three independent experiments. C) Representative immunofluorescence analysis of vGPCR-transfected human MSCs stained with SNA and PHA-L; fluorescence intensity displayed using fire-scale pseudocolor. D) Flow cytometry analysis of Gal-1, Gal-2, and Gal-7 binding in vGPCR-transfected human MSCs compared with GFP-empty vector control (N=4). E) Glycophenotypic profiling of human MSCs infected with rKSHV.219 using SNA, LEL, and PHA-L assessed by flow cytometry. Box plots represent rMFI from three independent experiments. F) Gal-1 binding to KSHV-infected and non-infected hMSCs. Left: representative histogram. Right: quantitative rMFI analysis of three independent experiments. G) Flow cytometry of SNA, PHA-L, and Gal-1 binding in human MSCs following stimulation with recombinant human (rh)-IL-6 (5 ng/ml), VEGFa (10 ng/ml), PDGF-a (10 ng/ml), PDGF-b (20 ng/ml), or FGF2 (50 ng/ml). Bars represent the mean ± SEM of three independent experiments. **P < 0.01, *P < 0.05.

Expression of vGPCR in hMSCs induced marked remodeling of this Gal-1 permissive profile, with reduced α(2-6) sialylation and increased β(1-6) N-glycan branching (Figure 2C and S2C). Strikingly, while Gal-1 binding was significantly increased under this condition, Gal-2 and Gal-7 binding remained unchanged (Figure 2D), indicating that vGPCR-driven glycosylation remodeling selectively enhances Gal-1 engagement, and implicating reduced α(2-6) sialylation as a key determinant of this selectivity.

To determine whether intact viral infection recapitulates these changes, hMSCs were infected with rKSHV.219 for 48 hours. KSHV-infected hMSCs exhibited decreased α(2-6) sialylation and increased β(1,6) N-glycan branching (Figure 2E), leading to a significant increase in Gal-1 binding (Figure 2F), indicating that virus-driven remodeling is sufficient to enhance Gal-1 association to hMSCs.

Finally, given that paracrine signaling has been proposed as a key contributor to KSHV-driven oncogenesis, we evaluated whether cytokines commonly associated with this process (IL-6, VEGF, PDGFs, and FGF2), could reproduce the observed glycosylation changes. None of these cytokines significantly altered SNA or PHA-L lectin binding profiles or modulated Gal-1 binding to hMSCs (Figure 2G), suggesting that glycan remodeling and selective Gal-1 binding reflects a virus-intrinsic mechanism rather than a secondary response to paracrine cytokine exposure.

### Glycosylation-related transcriptional programs are altered in KSHV infection and Kaposi sarcoma

To explore the molecular basis underlying the glycosylation changes observed *in vitro* upon infection and assess their relevance in human disease, we analyzed glycosylation-related gene expression profiles in two complementary transcriptomic datasets: KSHV-infected human MSCs^28^ and an integrative KS cohort comprising 131 samples, including control skin and KS tumors stratified into molecularly defined clusters, derived from multiple independent studies ^34^. Gene expression analysis was anchored in a curated set of carbohydrate-related genes compiled from established glycosylation pathway atlases^35,36^.

Analysis of the KSHV-infected hMSC model revealed significant dysregulation of pathways related to N-glycan biosynthesis, nucleotide-sugar transport, and carbohydrate metabolism (Figure S3A and S3B). Applying the same gene set to the integrative KS cohort, we identified 69 carbohydrate-associated genes differentially expressed between control skin and KS tumors (Figure 3A). Notably, cluster 1, as defined in the integrative KS cohort^34^ and characterized by enriched endothelial signatures and advanced tumor features, exhibited increased glycosylation-related gene expression. This observation suggests that glycosylation remodeling is not only a feature of KSHV infection *in vitro* but also a transcriptional hallmark associated with KS progression and molecular subtype.

**Figure 3.**
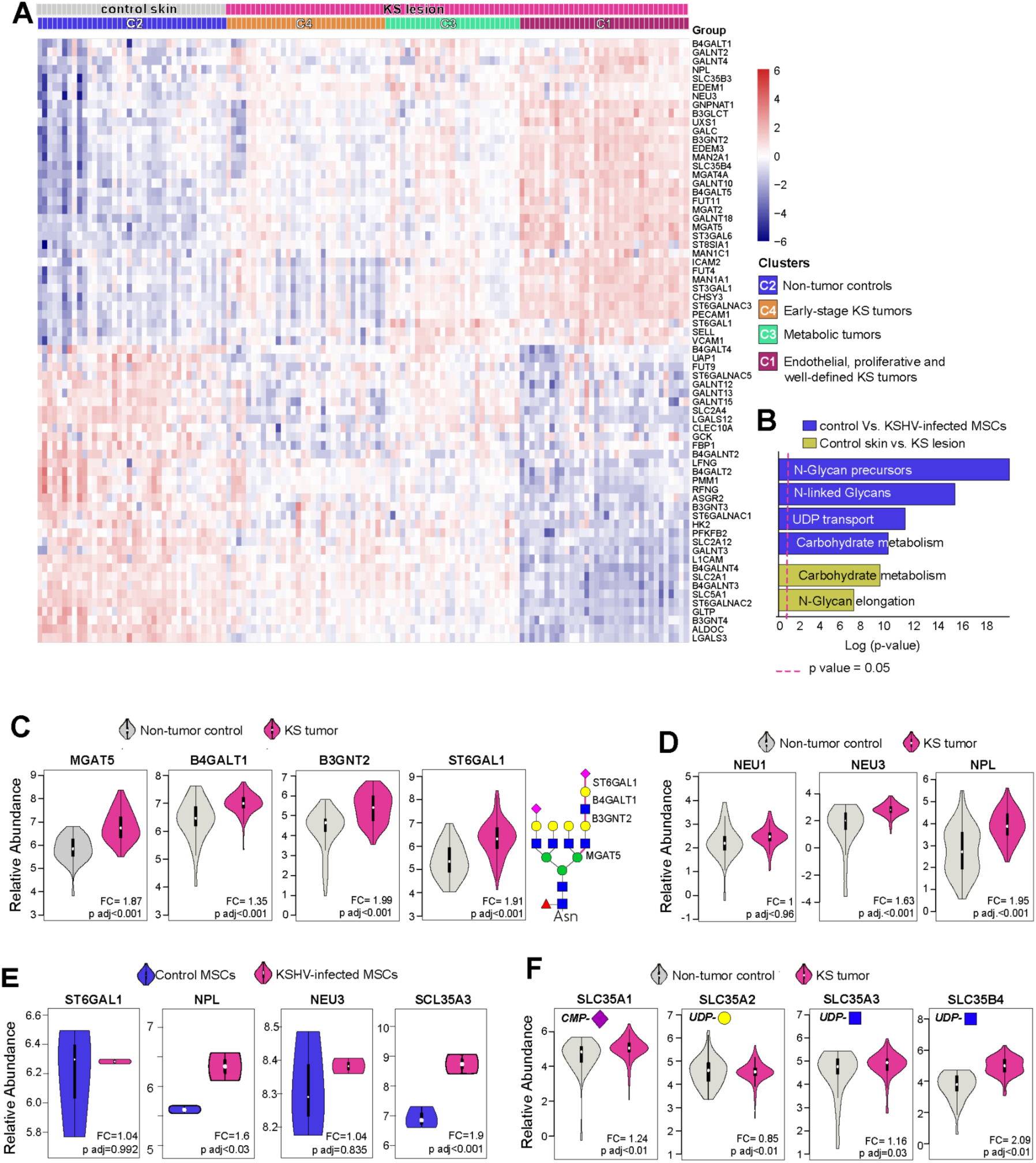
Transcriptional remodeling of glycosylation-related pathways in KSHV-infected MSCs and Kaposi sarcoma. A) Unsupervised hierarchical clustering of glycosylation-related gene expression in transcriptomic datasets derived from normal skin and Kaposi sarcoma (KS) lesions, stratified into molecularly defined tumor clusters. Gene set includes glycosyltransferases, glycosidases, nucleotide-sugar transporters, and lectins curated from established glycosylation pathway atlases. B) Gene Ontology enrichment analysis of glycosylation-related pathways in KS lesions versus normal skin and in KSHV-infected MSCs versus uninfected controls. C) Differential expression of genes involved in N-glycan branching and poly-LacNAc elongation, including MGAT5, B4GALT1, B3GNT2 and ST6GAL1, in KS lesions relative to normal skin. left panel, illustration of representative complex-N-glycan structure indicating the enzymes involved. D) Differential expression of genes involved in sialic acid metabolism and turnover, including NEU1, NEU3, and NPL, in KS lesions relative to normal skin. E) Transcriptomic analysis of ST6GAL1, NPL, NEU3 and SLC35A3 expression in KSHV-infected human MSCs compared with uninfected controls. F) Differential expression of Golgi-localized nucleotide-sugar transporter genes, including SLC35A1, SLC35A2, SLC35A3, and SLC35B4, in KS lesions relative to normal skin.

Gene Ontology enrichment analysis of KSHV-infected MSCs highlighted N-glycan biosynthesis, nucleotide-sugar transport, and carbohydrate metabolism as the most significantly altered pathways (Figure 3B). To further connect these *in vitro* observations with patient-derived data and identify relevant molecular nodes, we focused on three interconnected processes directly relevant to Gal-1–mediated receptor engagement: complex N-glycan formation, sialylation remodeling, and nucleotide-sugar transport, enzymatic axes that collectively determine Gal-1 ligand availability (Figure S3C).

Transcriptomic profiling revealed coordinated upregulation of genes involved in N-glycan branching and poly-LacNAc extension, including MGAT2, MGAT5, B3GNT2, B4GALT1, and B4GALT5 (Figure 3C and S3D). These enzymes collectively drive the formation of multi-antennary N-glycans with extended poly-LacNAc chains, structures known to serve as high-affinity ligands for Gal-1.

Lectin-binding assays in KSHV-infected cells revealed reduced α(2-6)–linked sialylation (Figure 2E); however, transcriptomic analysis of KS tumors demonstrated upregulation of the α(2,6) sialyltransferase 1 (ST6GAL1) (Figure 3C). To explore possible mechanisms that might reconcile this observation, we examined genes involved in sialic acid turnover. Neuraminidases NEU1 and NEU4 did not show consistent differential expression between KS and control samples (Figure 3D and S3C). In contrast, KS tumors showed upregulation of NEU3 (an enzyme that preferentially cleaves sialic acid from glycolipids), and N-acetylneuraminate lyase (NPL), enzyme that cleaves free sialic acid into pyruvate and N-acetylmannosamine, reducing glycan capping via decreased synthesis of CMP-sialic acid (Figure 3D). Additional dysregulation of sialyltransferases acting on lipids and O-glycans was also observed (Figure S3C), suggesting that in established tumors, sialylation is not simply reduced but actively redistributed through a combination of increased turnover and sustained sialyltransferase expression. In contrast, transcriptomic analysis of KSHV-infected hMSCs showed no significant changes in ST6GAL1 or neuraminidase expression but revealed selective upregulation of NPL (Figure 3E and S3A), suggesting that reduced sialylation in early infection may reflect altered sialic acid catabolism rather than transcriptional repression of sialyltransferases.

Finally, we examined Golgi-localized nucleotide-sugar transporters that supply substrates required for glycan biosynthesis. Transcriptomic profiling revealed increased expression of SLC35A1, SLC35A2 and SLC35A3/SLC35B4 in KS tumors (Figure 3F and S3C), which mediate Golgi import of CMP-sialic acid, UDP-Gal, and UDP-GlcNAc respectively.

Together, these data highlight three glycosylation-related modules associated with KS: N-glycan branching and poly-LacNAc extension (MGAT5/B3GNT2/B4GALT1), sialic acid turnover (NEU3/NPL), and nucleotide-sugar transport (SLC35A3/SLC35B4) (Figure S3D). Notably, the NPL/SLC35A3 axis was also upregulated in KSHV-infected MSCs, indicating that core features of the tumor glycosylation program are already engaged at early stages of KSHV infection.

### Galectin-1 couples glycosylation remodeling to PDGFRA activation and trafficking

Having established that KSHV infection and vGPCR expression remodel the mesenchymal cell glycome to favor Gal-1 binding, we next investigated whether enhanced lectin engagement has functional consequences for PDGFRA signaling.

First, we confirmed cell surface expression of both PDGFRA and PDGFRB in hMSCs by flow cytometry and confocal microscopy (Figure 4A). To test whether Gal-1 engagement modulates PDGFRA, hMSCs were incubated with recombinant Gal-1, and receptor phosphorylation was monitored over time. Gal-1 induced a clear increase in PDGFRA activation within 15 min of incubation (Figure 4B) and fluorescently labeled Gal-1 colocalized with phospho-PDGFRA at the cell surface (Figure S4A), indicating spatial proximity between Gal-1 and the activated receptor. Consistent with PDGFRA activation, Gal-1 stimulation also induced marked phosphorylation of STAT3, a canonical downstream effector of PDGFRA signaling (Figure 4C).

**Figure 4.**
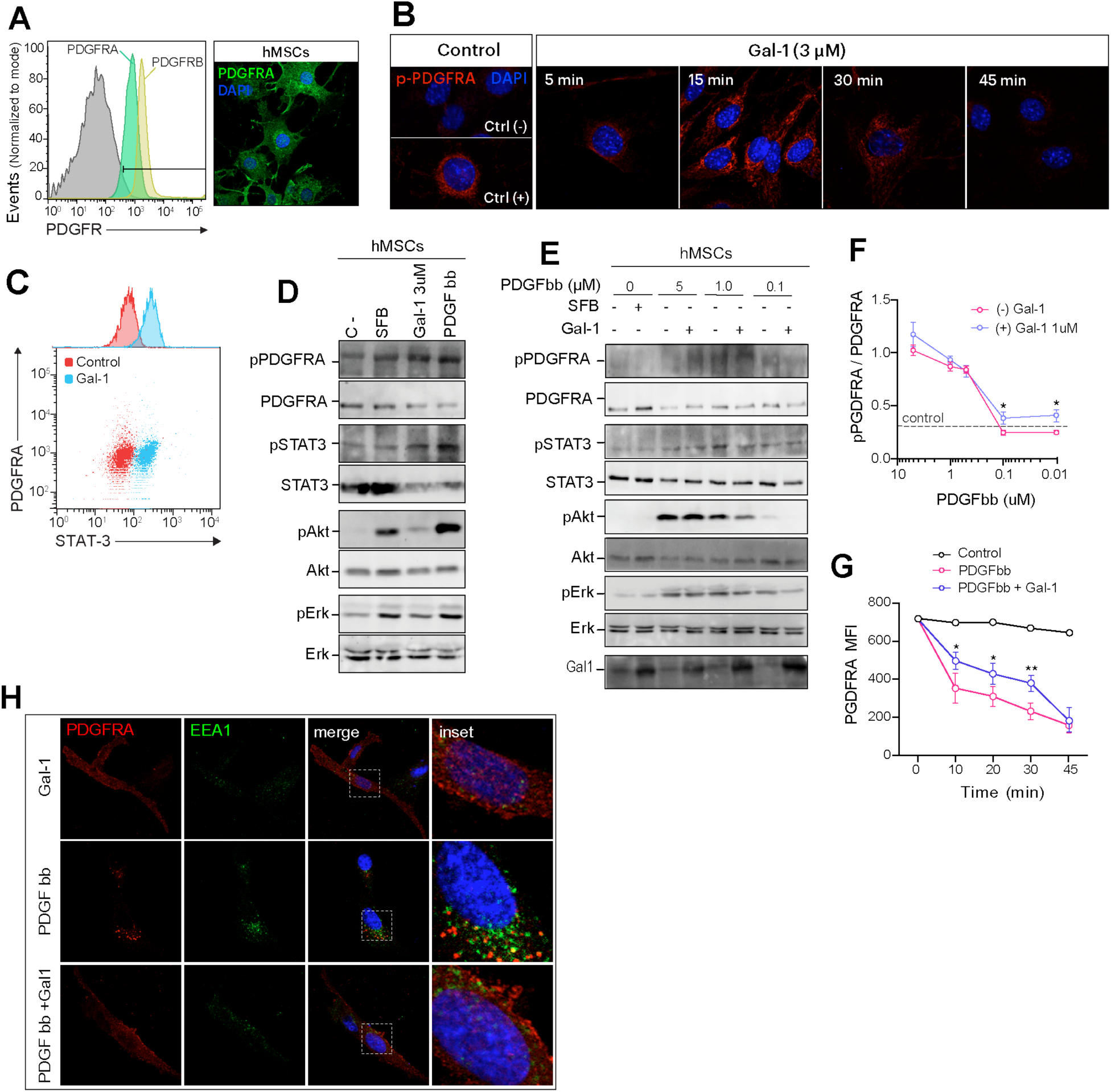
Gal-1al-1 directly activates PDGFRA signaling, sensitizes it to PDGF-BB, and delays its endocytic trafficking in human MSCs. A) Left: Flow cytometry analysis of PDGFRA and PDGFRB surface expression in human MSCs. Right: confocal microscopy image of PDGFRA distribution. B) Immunofluorescence analysis of phosphorylated PDGFRA at indicated time points following incubation with Gal-1 (3μM). Negative control: serum-free medium; positive control: PDGF-BB (5μM). C) Flow cytometry analysis of STAT3 phosphorylation in human MSCs treated with Gal-1 (3 μM). D) Immunoblot analysis of PDGFRA, STAT3, AKT, and ERK1/2 phosphorylation in human MSCs under serum-free, serum-containing, recombinant Gal-1 (3 μM), and PDGF-BB (5μM) conditions. E) Immunoblot analysis of PDGFRA, STAT3, AKT, and ERK1/2 phosphorylation in MSCs co-stimulated with recombinant Gal-1 (1 μM) and decreasing concentrations of PDGF-BB. F) Quantification of PDGFRA phosphorylation dose-response shown in E, expressed as the ratio of phosphorylated to total PDGFRA (N=4). G) Flow cytometry analysis of PDGFRA surface levels over time in hMSCs treated with PDGF-BB alone or PDGF-BB + Gal-1. (N=4). H) Confocal microscopy of PDGFRA and EEA1 in hMSCs treated with Gal-1, PDGF-BB, or PDGF-BB + Gal-1 at 30 min. Images are representative of at least three independent experiments. Immunoblots and flow cytometry data are representative of three independent experiments. Bars represent mean ± SEM. **P < 0.01, *P < 0.05.

These observations were corroborated by immunoblot analysis, which demonstrated that Gal-1 induced phosphorylation of PDGFRA and its downstream effectors, AKT and ERK1/2, in hMSCs (Figure 4D). This signaling response was conserved in mMSCs and the mesenchymal cell line OP9 (Figure S4B-D) following Gal1-stimulation. Although the magnitude of Gal-1–induced activation was lower than that elicited by PDGF-BB, the pattern of pathway engagement was qualitatively comparable, consistent with Gal-1’s role as a non-canonical PDGFRA activator rather than a full agonist.

Beyond direct receptor activation, galectins are known to modulate receptor responsiveness to canonical ligands by stabilizing receptor complexes at the cell surface ^11,39^. To test whether Gal-1 sensitizes PDGFRA to its canonical ligand, hMSCs were co-stimulated with recombinant Gal-1 and decreasing concentrations of PDGF-BB. PDGFRA phosphorylation was detectable at significantly lower PDGF-BB concentrations in the presence of Gal-1 than when as compared to cells exposed to PDGF-BB alone (Figure 4E and 4F), suggesting that Gal-1 lowers the ligand threshold required for receptor activation. This data suggests that a Gal-1–permissive glycan environment could sustain PDGFRA signaling even under limiting ligand availability.

To determine whether Gal-1 influences PDGFRA receptor dynamics, we assessed PDGFRA internalization kinetics by monitoring cell surface PDGFRA levels over time. Stimulation with PDGF-BB induced the expected progressive reduction in surface PDGFRA, consistent with ligand-driven RTK internalization (Figure S4E). In contrast, simultaneous exposure of PDG-BB and Gal-1 resulted in significantly higher levels of PDGFRA retained at the cell surface across early time points (Figure 4G), indicating that Gal-1 engagement delays receptor endocytosis, extending the window of receptor availability at the plasma membrane.

To visualize these differences at the subcellular level, we examined PDGFRA localization relative to the early endosomal marker EEA1 by confocal microscopy. In control cells and in cells stimulated with PDGF-BB, PDGFRA displayed prominent intracellular localization and partial co-localization with EEA1, consistent with active receptor trafficking to early endosomes (Figure 4H). In contrast, cells exposed to Gal-1 retained substantial PDGFRA at the plasma membrane with reduced EEA1 co-localization, indicating that Gal-1 engagement stabilizes the receptor at the cell surface and restricts its endocytic trafficking.

Together, these results indicate that Gal-1 enhances PDGFRA signaling through a dual mechanism: direct receptor activation and lattice-dependent retention at the plasma membrane, both of which are enabled by the Gal-1–permissive glycan landscape generated by KSHV infection in mesenchymal stromal cells.

## Discussion

Successful viral infection demands more than subversion of signaling pathways: it requires the capacity to operate within, and selectively reshape, the regulatory architecture of the host cell without dismantling it ^3,40^. One of such layers is protein glycosylation. The glycan landscape of the cell surface controls receptor organization, membrane retention, and activation thresholds independently of receptor abundance or ligand availability^10,13,14,41^. Oncogenic herpesviruses are known to exploit these layers, consistent with the selective, coordinated nature of virus-driven glycosylation remodeling documented in infected cells^42^. Whether this remodeling feeds back onto receptor signaling competence remains unexplored.

The data presented here establish that KSHV reprograms the glycosylation landscape of mesenchymal stromal cells through coordinated metabolic, enzymatic, and transport-level adaptations, generating a cell surface environment selectively enriched in Gal-1 permissive glycan structures. This remodeled glycome shapes receptor tyrosine kinase signaling through three convergent mechanisms: Gal-1–dependent PDGFRA activation, reduced ligand threshold, and delayed endocytic trafficking. The functional relevance of this sensitization effect is amplified in the context of KSHV pathogenesis, where paracrine PDGF concentrations may be suboptimal for full receptor activation. Glycosylation remodeling, in this context, emerges as an active determinant of signaling competence in KSHV-conditioned mesenchymal cells.

The oncogenic role of PDGFRA signaling in KS is well established, particularly in the context of KSHV-driven autocrine and paracrine signaling loops that sustain mesenchymal and endothelial cell activation ^5–7,43^. vGPCR, as a constitutively active signaling hub reinforce proliferative, converges on PDGFRA-associated pathways to, angiogenic, and survival programs central to lesion maintenance ^23,44,45^. Our findings highlight glycosylation remodeling as an additional regulatory dimension through which KSHV shapes the signaling competence of mesenchymal cells. This interpretation gains further support from the observation that N-linked glycosylation is required for vGPCR trafficking and oncogenic activity^22^, suggesting that glycan-dependent regulation operates at multiple levels of the KSHV oncogenic program, from viral protein function to host receptor signaling.

The regulation of RTK signaling through lectin–glycan interactions has emerged as a central principle in cancer biology ^20,24,25,27,46^, yet its intersection with viral oncogenesis has not been explored. Oncogenic viruses, including herpesviruses, are known to alter host glycosylation ^42,47^. Notably, systematic glycoproteomic profiling of herpesvirus-infected cells has revealed this remodeling is selective and coordinated, engaging specific components of the glycosylation machinery^42^, a principle consistent with this pathway-specific transcriptional changes we observe in KSHV-infected MSCs and KS tumors and with the functional consequences for RTK signaling documented here.

Within this context, Gal-1 emerges not only as a soluble effector of the tumor microenvironment but also as a contextual sensor of virus-induced glycan remodeling in which functional output is determined by the glycan landscape it encounters as well as its own expression level. Gal-1 binding is dictated primarily by the density, branching, and terminal modification of LacNAc-containing glycans ^21,48,49^. Our data indicate that KSHV infection and vGPCR signaling operate on both sides of this equation simultaneously: increasing Gal-1 expression^16^ while reshaping the cell surface glycome to favor lectin engagement through enhanced N-glycan branching and reduced α(2-6)–sialylation. The functional consequence is the formation of Gal-1–glycan lattices at the cell surface, supramolecular assemblies known to stabilize receptors in signaling-competent membrane domains, reduce endocytic turnover, and lower activation threshold^39,50,51^. Our observation that Gal-1 delays PDGFRA internalization and sustains receptor phosphorylation provides direct functional evidence for this mechanism in the context of KSHV-conditioned mesenchymal cells.

Beyond changes in glycan branching, our transcriptomic analyses point to coordinated remodeling of sialic acid metabolism as a defining feature of KSHV-associated glycosylation changes. The reduction in α(2-6) sialylation observed in KSHV-infected MSCs and KS samples does not appear to result from decreased ST6GAL1 expression or increased neuraminidase expression as described in other oncogenic contexts^52,53^. Instead, we observed consistent upregulation of NPL, an enzyme that cleaves sialic acid into pyruvate and N-acetylmannosamine^54,55^. This observation suggests that KSHV may redirect sialic acid away from terminal glycan capping toward metabolic reutilization. Unlike neuraminidase-mediated desialylation, which releases free sialic acid for recycling, NPL-driven catabolism produces pyruvate and ManNAc, intermediates that can feed into glycolytic and hexosamine biosynthetic pathways respectively ^54,55^, potentially linking glycan remodeling to broader metabolic reprogramming in infected cells.

The selectivity of this remodeling effect extends to the transport level. Among the Golgi-localized nucleotide-sugar transporters examined, those supplying substrates for N-glycans branching and sialylation appear consistently upregulated such as SLC35A3, SLC35B4 and SLC35A1 with mediate import of UDP-GlcNAc, UDP-GalNAc and CMP-sialic acid respectively^56^. Transporters governing other glycosylation pathways were largely unchanged. This substrate-level specificity confirms that KSHV-driven glycosylation remodeling is a directed program.

Poly-LacNAc elongation results from the alternating activity of β(1-3) N-acetylglucosaminyltransferases and β(1-4) galactosyltransferases, most prominently B3GNT2 and B4GALT1, which together generate linear repeats of the Galβ1-4GlcNAc (LacNAc) disaccharide on branched N-glycans^57^. These extended LacNAc chains constitute canonical ligands for galectins and are critical determinants of lectin–glycan lattice formation at the cell surface^11^. The coordinated upregulation of MGAT5 together with B3GNT2 and B4GALT1, points to selective activation of the glycosylation machinery to generate a glycan architecture optimized for Gal-1 engagement.

Functionally, these glycosylation changes have direct consequences for PDGFRA signaling dynamics. Our data indicate that Gal-1 promotes PDGFRA activation and lowers the activation threshold of this receptor in response to PDGF-BB, a phenomenon consistent with lattice-dependent receptor priming described for other RTK systems ^11,20,39,46,58,59^. Mechanistically, Gal-1 also delays PDGFRA internalization and reduces its trafficking to early endosomes, extending the residence time of the activated receptor at the plasma membrane and thereby prolonging downstream signaling^39,46,50^. This mechanism may operate both in lytically infected vGPCR-expressing cells and in bystander uninfected cells exposed to secreted Gal-1 and paracrine PDGF, a combination that, in a remodeled glycan environment, could sustain oncogenic signaling beyond the infected cell compartment.

Taken together, our findings establish a coordinated and selective reprogramming of mesenchymal cell glycosylation through metabolic, enzymatic, and transport-level adaptations in KSHV-infected cells and KS tumors. These changes generate a cell surface environment that favors Gal-1 cross-linking and sustains PDGFRA signaling, not as independent oncogenic drivers, but as permissive adaptations that lower the threshold for signaling events already central to KS pathogenesis. Thus, KSHV exploits host this glycosylation-dependent regulatory circuit to persist within mesenchymal tissue. Sarcomagenesis is the cost the host pays. Glycome remodeling operates through defined molecular nodes, arising through transcriptional signatures traceable in patient-derived data, and has direct functional consequences for RTK activation. Whether analogous programs operate in other herpesvirus-associated malignancies, and whether the NPL/SLC35A3 axis represents a shared strategy among oncogenic herpesviruses, are questions that warrant further exploration.

## Lead contact

Further information and requests for resources should be directed to and will be fulfilled by the lead contact, Diego O. Croci.

## Materials availability

This study did not generate new unique reagents.

## Acknowledgments

The work was supported by NIH Award Number U54CA221208 (GAR and DOC), CONICET (PIP2022) to DOC, Argentinean Agency of Science, Technology and Development (PICT2019-0532 and PICT2014-0205) to DOC. We thank support from the SALES Foundation, and Ferioli-Ostry family. We would also like to thank Omar Coso for support and continuous advice, Huésped Foundation team Buenos Aires (Pedron Cahn, Omar Sued and Valeria Fink) for intellectual support and Kerry Burstein for intellectual advice. We especially thank the U54-consortium for Research and Training in Virally Induced AIDS-Malignancies UM-Argentina. We dedicate this article to Enrique Mesri, the driving force and coordinator of this consortium, who sadly passed during the development of this project.

## Author Contributions

Experimental design J.G.T., J.P.C., J.N., D.O.C., Data analysis and interpretation J.G.T., P.N.G, B.N., P.A.G., K.M., J.N., E.L., D.O.C.; KSHV infection experiments M.A.M., J.N., P.N.G.;

Experimental execution J.G.T., P.N.G., M.A.M., B.N., P.A.G., J.N.; Bioinformatic analyses M.A., J.N., E.L., P.A.G.; Conceptual design D.O.C., J.P.C, E.A.M., G.A.R.; Writing – original draft,

D.O.C. and J.G.T.; review & editing, all the authors.

## Declaration of AI-assisted technologies in the writing process

During the preparation of this work the authors used Claude Sonnet 4.6 to improve the readability and language of the manuscript. After that each author reviewed and edited the content as needed and takes full responsibility for the content of the published article.

