## supplemental figures for "Kaposi Sarcoma Herpes Virus Reprograms Mesenchymal Cell Glycosylation to control platelet-derived growth factor receptor A signaling": Gambarte Tudela et al Supplementary Data.pdf

†Passed, August 27, 2022

**Graphical Abstract.** KSHV infection remodels the mesenchymal cell glycome through coordinated regulation of N-glycan branching, sialic acid metabolism, and nucleotide-sugar transport. These changes ultimately assemble Gal-1–permissive glycan structures that promote lectin-dependent PDGFRA signaling and contribute to KS pathogenesis.

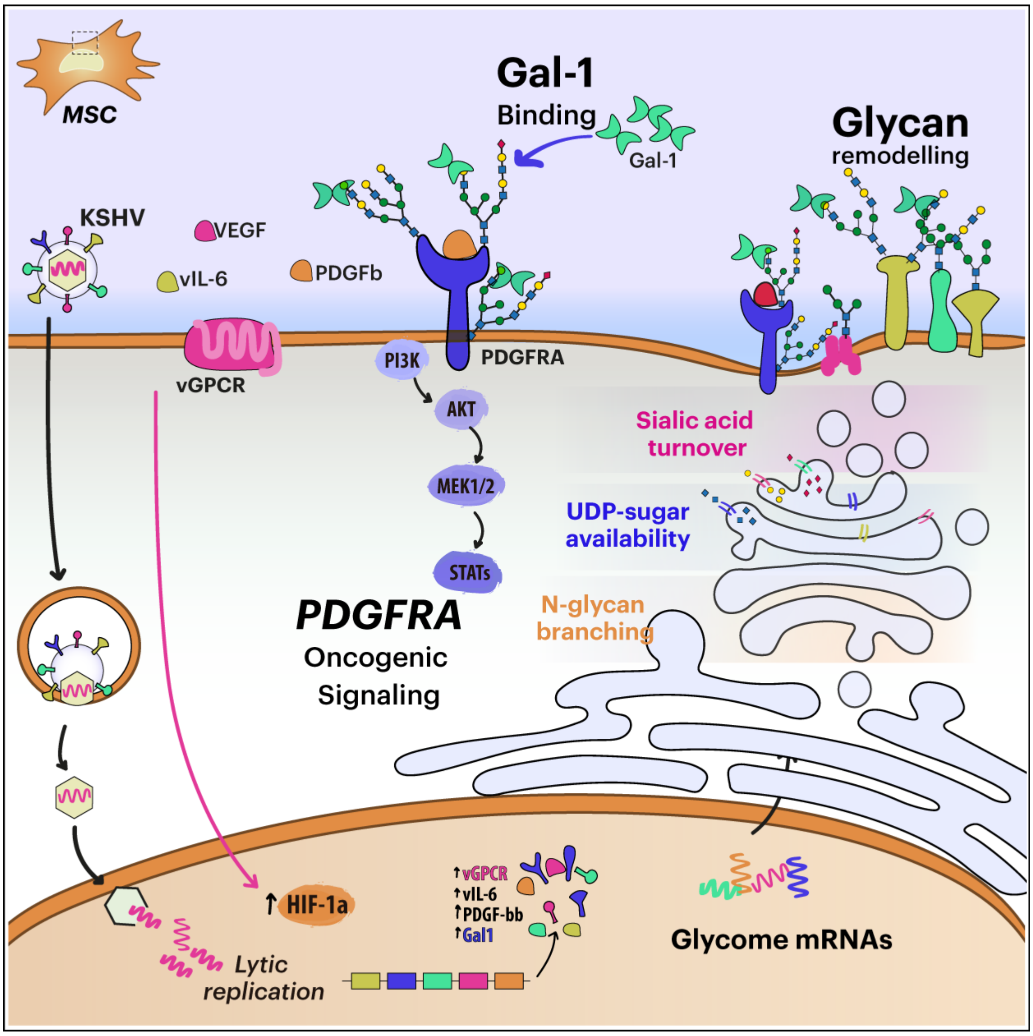

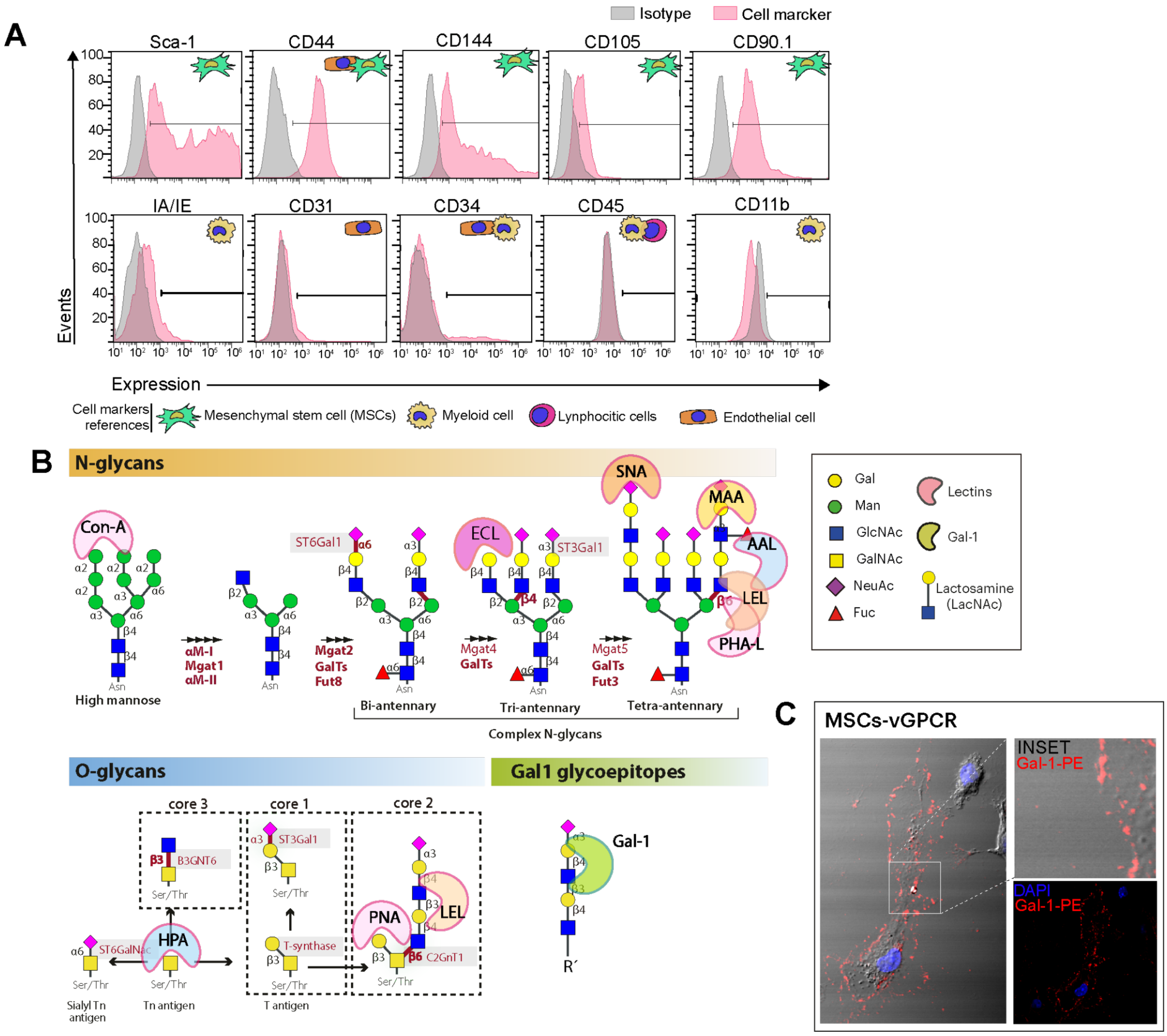


**Figure S1. Phenotypic characterization of murine BM-MSCs, lectin panel reference, and Gal-1 binding in vGPCR-expressing cells.** A) Flow cytometry analysis of mouse MSC surface phenotype. Cells were characterized using positive markers and negative markers confirming MSC identity and ruling out contamination with myeloid, lymphocytic, and endothelial cell populations. B) Schematic reference panel illustrating the glycan structures recognized by lectins and Gal-1 used in this study. Upper panel: N-glycan biosynthetic pathway showing high-mannose, bi-antennary, tri-antennary, and tetra-antennary complex N-glycans, with the glycosyltransferases responsible for each branching step and the lectins that recognize each structure (Con-A, ECL, SNA, MAA, AAL, LEL, PHA-L). Lower left panel: O-glycan core structures (core 1, core 2, core 3) and their biosynthetic relationships, with lectins HPA and PNA indicated. Lower right panel: schematic of the minimal Gal-1 binding glycoepitope, a poly-LacNAc unit on branched N- or O-glycans. C) Confocal microscopy of Gal-1 binding in vGPCR-expressing BM-MSCs. The main panel shows Gal-1–PE staining at the cell surface; inset shows higher magnification with DAPI counterstain. Images are representative of three independent experiments

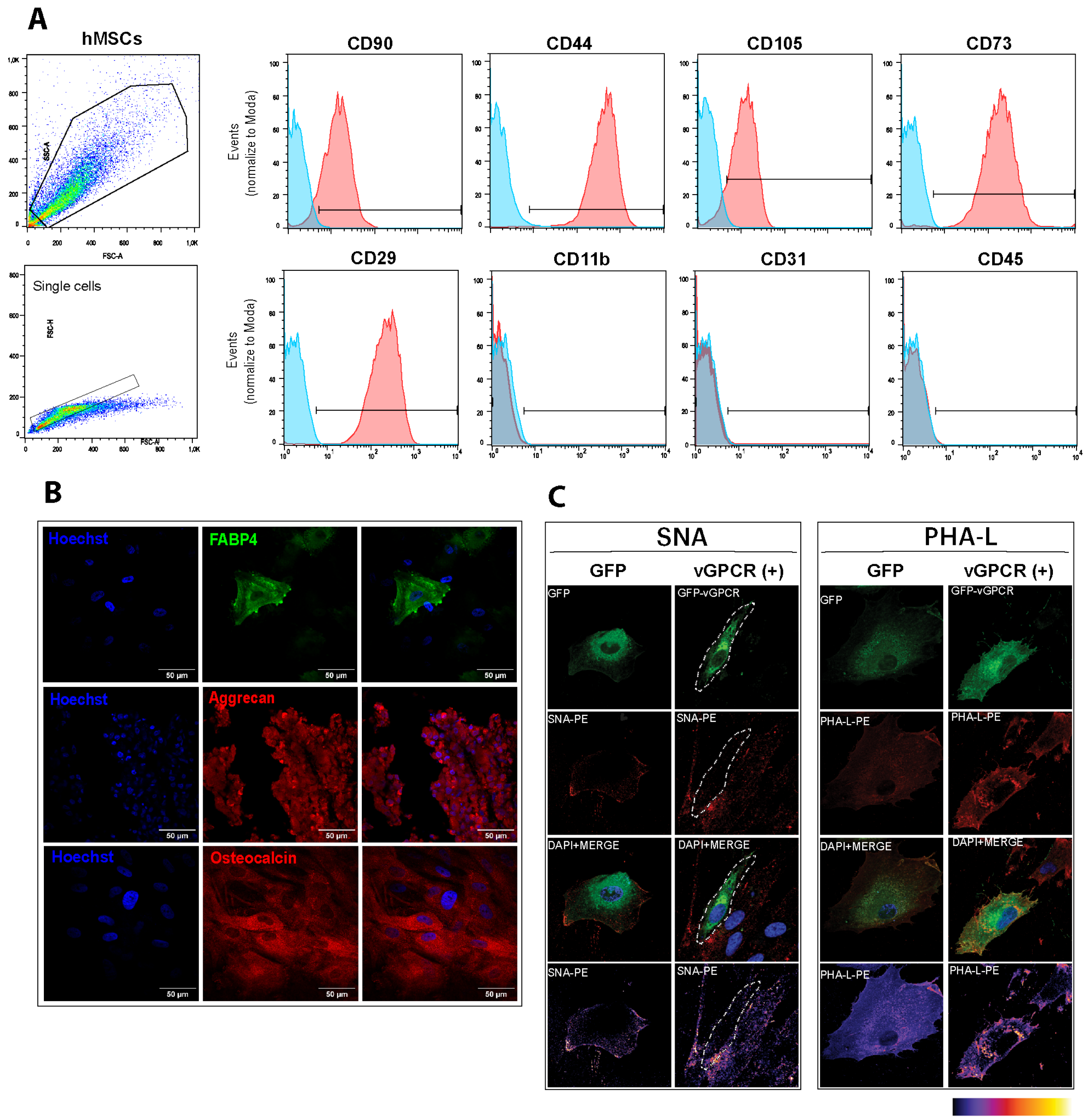


**Supplementary Figure 2. Characterization of human umbilical cord–derived mesenchymal stromal cells and glycosylation changes induced by vGPCR expression.** A) Flow cytometry analysis of human MSC surface phenotype. Cells were characterized using positive markers CD90, CD44, CD105, CD73, and CD29, and negative markers CD11b, CD31, and CD45, confirming MSC identity according to ISCT criteria. B) Multilineage differentiation capacity of hMSCs. Cells were induced toward adipogenic (FABP4), chondrogenic (Aggrecan), and osteogenic (Osteocalcin) lineages and assessed by immunofluorescence. C) Representative confocal microscopy images of SNA and L-PHA binding in human MSCs transfected with vGPCR-GFP or GFP-empty vector. Fluorescence intensity displayed using fire-scale pseudocolor. Images are representative of three independent experiments.

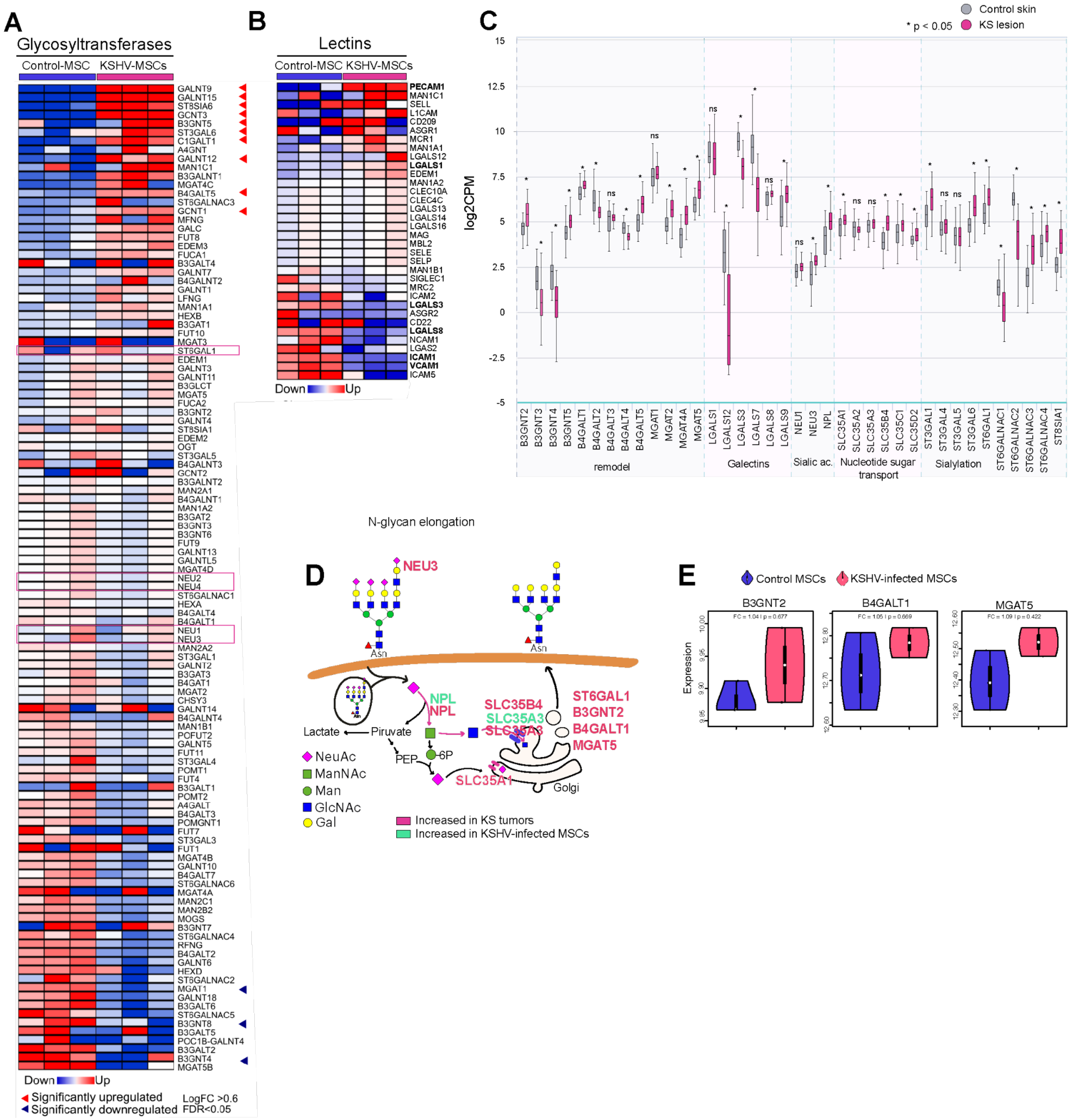


**Figure S3. Extended transcriptomic analysis of glycosylation-related gene expression in KSHV-infected MSCs and Kaposi sarcoma.** A) Unsupervised hierarchical clustering of glycosyltransferase gene expression in control and KSHV-infected human MSCs. Red arrowheads indicate significantly upregulated genes (LogFC > 0.6, FDR < 0.05); blue arrowheads indicate significantly downregulated genes. B) Unsupervised hierarchical clustering of lectin gene expression in control and KSHV-infected MSCs. Boldface gene names indicate statistically significant differential expression. C) Differential expression analysis of glycosylation-related genes in KS lesions compared with control skin, grouped by functional category: N-glycan elongation, galectins, sialic acid remodeling, nucleotide-sugar transport, and sialylation. Data expressed as log2CPM. *$p < 0.05$; ns, not significant. D) Schematic representation of the glycosylation remodeling program identified in

KSHV-infected MSCs and KS tumors, highlighting the NPL/SLC35A3 metabolic axis and its relationship to N-glycan branching and sialic acid catabolism. Pink labels indicate genes upregulated in KS tumors; green labels indicate genes upregulated in KSHV-infected MSCs. E) Violin plots of B3GNT2, B4GALT1, and MGAT5 expression in control MSCs versus KSHV-infected MSCs. FC and adjusted p-values are indicated.

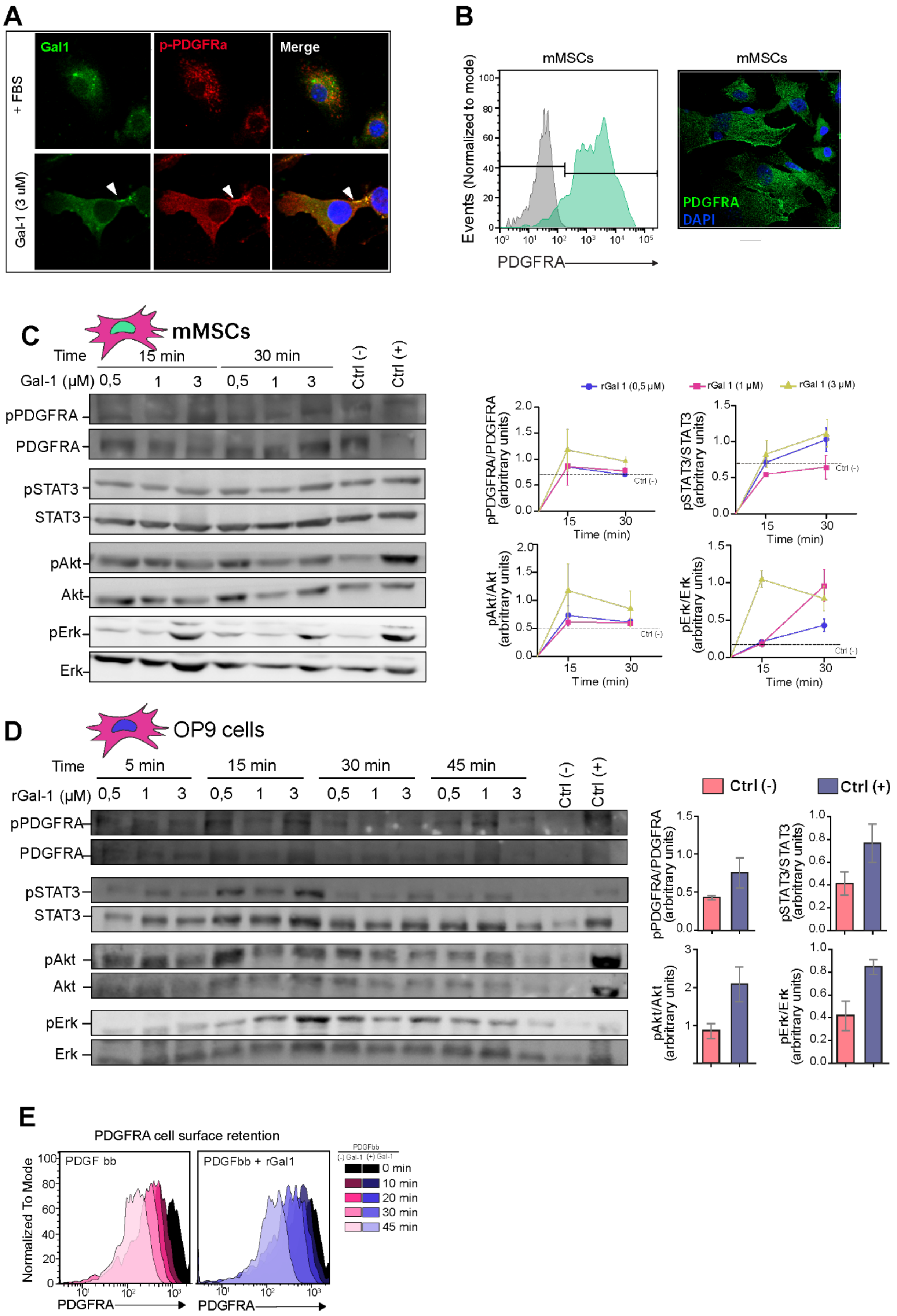
A
Gal1
p-PDGFRa
Merge
+ FBS
Gal-1 (3 uM)
B
mMSCs
Events (Normalized to mode)
PDGFRA
PDGFRA
DAPI
C
mMSCs
Time
15 min
30 min
Gal-1 (μM)
Ctrl (-)
Ctrl (+)
pPDGFRA
PDGFRA
pSTAT3
STAT3
pAkt
Akt
pErk
Erk
rGal 1 (0,5 μM)
rGal 1 (1 μM)
rGal 1 (3 μM)
pPDGFRA/PDGFRA (arbitrary units)
pSTAT3/STAT3 (arbitrary units)
pAkt/Akt (arbitrary units)
pErk/Erk (arbitrary units)
Time (min)
Ctrl (-)
D
OP9 cells
Time
5 min
15 min
30 min
45 min
rGal-1 (μM)
Ctrl (-)
Ctrl (+)
E
PDGFRA cell surface retention
PDGF bb
PDGFbb + rGal1
Normalized To Mode
PDGFRA
0 min
10 min
20 min
30 min
45 min

**Figure S4. Gal-1–driven PDGFRA activation and surface retention in murine mesenchymal cells and OP9 cells.** A) Confocal microscopy of Gal-1 and phospho-PDGFRA co-localization in human MSCs treated with recombinant Gal-1 (3 μM). Arrowheads indicate co-localization of Gal-1 and p-PDGFRA at the plasma membrane. B) Left: flow cytometry analysis of PDGFRA surface expression in murine mMSCs. Right: confocal microscopy image of PDGFRA in mMSCs; nuclei counterstained with DAPI. C) Left: immunoblot analysis of PDGFRA, STAT3, AKT, and ERK1/2 phosphorylation in mMSCs treated with recombinant Gal-1 (0.5, 1, or 3 μM) for 15 or 30 min. Right: quantification of phosphorylation ratios over time for each Gal-1 concentration. D) Left: immunoblot analysis of PDGFRA, STAT3, AKT, and ERK1/2 phosphorylation in OP9 mesenchymal cells treated with recombinant Gal-1 (0.5, 1, or 3 μM) for 5, 15, 30, or 45 min. Right: quantification of phosphorylation ratios at Ctrl (−) and Ctrl (+) conditions. E) Flow cytometry analysis of PDGFRA cell surface retention over time (0, 10, 20, 30, 45 min) in human omaMSCs treated with PDGF-BB alone or PDGF-BB + recombinant Gal-1, showing delayed receptor internalization in the presence of Gal-1. Immunoblots and histograms are representative of at least three independent experiments. Bars represent mean ± SEM.
